# Microbiome-oriented data mining of operational monitoring of anaerobic digestion reactor during steady operation period, failure, and restoration

**DOI:** 10.1101/2025.02.07.637110

**Authors:** M. Panou, I. Kavakiotis, Anastasios Mitsopoulos, Vasiliki Tsioni, Themistoklis Sfetsas

**Affiliations:** QLAB Private Company. Research & Development, Quality Control and Testing Services, 57008 Thessaloniki, Greece; Toolboks LP, Athens, Greece; Ergoplanning Ltd. Consulting, 57300 Thessaloniki, Greece

## Abstract

Anaerobic digestion (AD) is an essential biotechnology for sustainable waste management and bioenergy production, driven by a complex microbial ecosystem. However, the stability and efficiency of AD reactors are frequently challenged by microbial shifts resulting from operational disturbances. This study employs microbiome-oriented data mining and bioinformatics approaches to analyze microbial community dynamics in an anaerobic digestion reactor under steady-state operation, failure, and restoration phases. Data were extracted from a biogas plant and used to evaluate a system failure, providing crucial insights into microbial shifts associated with operational instability. A custom-built microbiome-oriented database, μ2Gas, was developed to integrate sequencing data with physicochemical monitoring results, enabling structured data storage, retrieval, and correlation analysis. This database was instrumental in identifying key microbial taxa responsible for reactor destabilization and in predicting failure events based on microbial and physicochemical trends. Through high-throughput 16S rRNA gene sequencing and advanced bioinformatics analysis, microbial shifts were correlated with key operational parameters such as pH, volatile fatty acids (VFAs), and ammonia concentrations. Results indicate that methanogenic archaea, particularly Methanosarcina, experienced a sharp decline during reactor failure, followed by a gradual recovery, demonstrating their sensitivity to environmental stressors. Additionally, the balance between Firmicutes and Actinobacteria was identified as a crucial determinant of reactor stability, with Firmicutes recovering in tandem with system restoration. Feedstock composition was found to play a significant role in microbial shifts, with seasonal variability influencing community composition and reactor performance. Machine learning models applied to the database suggested the potential for predictive analytics in anticipating system failures. By leveraging microbial abundance patterns, these models were able to identify early warning signs of instability. Restoration efforts, including feedstock adjustments and operational parameter optimization, successfully reinstated core microbial communities essential for biogas production. This study underscores the critical role of microbiome monitoring in anaerobic digestion and highlights the potential of bioinformatics tools for enhancing reactor resilience. Future research should focus on real-time microbiome tracking, machine learning-driven predictive maintenance, and adaptive operational strategies to further improve AD efficiency and biogas yield.

## 1. Introduction

Anaerobic digestion (AD) is a crucial biotechnology for sustainable waste management and bioenergy production. It converts organic matter into biogas, primarily methane (CH4) and carbon dioxide (CO2), through a series of microbial-driven biochemical reactions (Angelidaki et al., 2011). The process is carried out by a highly diverse and functionally specialized microbial community that operates under strictly anaerobic conditions (Sundberg et al., 2013). Key microbial groups include hydrolytic, fermentative, acetogenic, and methanogenic microorganisms, which work sequentially to break down complex organic matter into methane (Ahring et al., 2003).

Despite its advantages, AD is highly sensitive to disturbances, such as variations in feedstock composition, accumulation of inhibitory compounds (e.g., ammonia, sulfides, volatile fatty acids), and fluctuations in operational parameters like pH and temperature (Chen et al., 2008). These disruptions can lead to microbial community shifts, process inhibition, and in extreme cases, reactor failure (Schnürer & Jarvis, 2010). Monitoring microbial community dynamics is crucial for understanding how these shifts affect reactor performance and resilience during steady-state operation, failure, and subsequent recovery (Ziganshin et al., 2013).

Advances in high-throughput sequencing, particularly 16S rRNA gene amplicon sequencing, have enabled comprehensive taxonomic and functional characterization of AD microbial communities (Callahan et al., 2016). Integrating microbiome analysis with physicochemical monitoring allows for more precise predictions of system instability and aids in developing strategies for maintaining reactor efficiency (Vanwonterghem et al., 2016). Despite these advancements, there remains a need for robust bioinformatics tools and databases to improve the interpretation of microbiome data in AD systems.

This study employs microbiome-oriented data mining and bioinformatics approaches to investigate the microbial composition and dynamics of an anaerobic digestion reactor across steady-state operation, failure, and restoration phases. By integrating microbiome data with key operational parameters, this work aims to identify critical microbial taxa and their functional roles in reactor stability and recovery. The findings contribute to the optimization of AD processes, enhancing resilience to disturbances and improving the sustainability of biogas production.

## 2. Materials and Methods

### 2.1 Determination of Physicochemical Parameters

Chemical oxygen demand (COD), total nitrogen (N), and total phosphorus (P) were measured for both influent synthetic wastewater and effluent samples using Hach LCK kits with a HACH DR 3900 spectrophotometer. These analyses were performed weekly. The concentrations of ammonium nitrogen (N-NH4), nitrate nitrogen (N-NO_3_), and nitrite nitrogen (N-NO_2_) were determined according to standard methodologies (Rice et al., 2017) using a JASCO V-630 spectrophotometer. The mixed liquor suspended solids (MLSS) and mixed liquor volatile suspended solids (MLVSS) were also quantified following standard methods for water and wastewater examination (Rice et al., 2017).

### 2.2 DNA extraction and 16S rRNA gene amplicon sequencing

Genomic DNA was extracted from the biofilm suspensions with the DNeasy PowerSoil Pro Kit (QIAGEN, Hilde, Germany) according to the manufacturer’s instructions. The quantity and quality of the extracted DNA were then estimated using a V-630 Spectrophotometer (JASCO, Inc, Japan). Library preparation was performed following the standard guidelines of the 16S Metagenomic Sequencing Library Preparation protocol (Illumina^TM^, Inc., San Diego, CA, United States). In brief, DNA was amplified using the HotStarTaq® Master Mix Kit (QIAGEN, Hilde, Germany) with the addition of the 341f/805r primer pair, which targets the bacterial and archaeal V3–V4 hypervariable regions of the 16S rRNA gene (341f 5′- CCTACGGGNGGCWGCAG-3′, 805r 5′-GACTACHVGGTATCTAATCC-3′). The PCR mixture (25 μL) contained 12.5 μL of HotStarTaq Master Mix, 5 μL of each primer, and 2.5 μL of DNA (5 ng/μL). Thermal cycling conditions included an initial 3-minute step at 95°C, followed by 25 cycles of denaturation at 95°C for 30 sec, annealing at 55°C for 30 sec and elongation at 72°C for 30 sec and a final extension step at 72°C for 5 min. PCR amplicons were cleaned up by AMPure XP beads (Beckman Coulter, CA, USA) to remove unbound primers and primer dimers. Next, dual indices and Illumina sequencing adaptors were attached with an index PCR using the Nextera XT Index Kit (Illumina Inc., San Diego, CA, USA). The PCR reaction mixture (50 μL) comprised 25 μL of HotStarTaq Master Mix, 5 μL of each index, 10 μL of PCR Grade Water and 5 μL of the previous PCR product, and the cycling conditions remained the same as that of the first PCR reaction except that the number of iterative cycles was reduced to 8. Afterwards, Indexed PCR amplicons were cleaned up using the AMPure XP beads (Beckman Coulter, CA, USA). The produced DNA libraries were quantified with the Qubit™ 4 Fluorometer (Thermo Fisher Scientific, Waltham, MA, USA), and their size was verified via a 1.5 % agarose gel electrophoresis. Equimolar concentrations of the libraries were then pooled together, and a quantitative PCR was performed using the QIAseq Library Quant Assay Kit (QIAGEN, Germany) for library concentration evaluation. The pooled library was subsequently spiked with 25% phiX control library (Illumina Inc., San Diego, CA, USA), denatured and diluted to a final concentration of 6 pM. Sequencing was performed on an Illumina MiSeq^TM^ platform with the MiSeq Reagent Nano Kit version 2 (500-Cycle)/ MiSeq Reagent Kit version 3 (600-Cycle) chemistry for a paired-end, 2×250-bp/2×300 cycle run.

### 2.3 Bioinformatics

BASESPACE. In BaseSpace, raw reads from each run were quality checked using FastQC (v.1.0.0, BaseSpace Illumina) and subsequently trimmed and quality filtered using FastqTool (v.2.2.5, BaseSpace Illumina). Genome *de novo* assembly and assembly quality were performed using SPAdes (v.3.9.0, BaseSpace Illumina). BaseSpace Bacterial Analysis Pipeline was used for species determination and antimicrobial resistance genes (ARGs) identification (v.1.0.4, BaseSpace Illumina). BaseSpace is equipped with a “Bacterial Analysis Pipeline” that claims to be able to predict the species of bacterial input genomes (using KmerFinder [<u>15</u>, <u>17</u>]), identifying ARGs (with ResFinder [<u>18</u>, <u>19</u>]), and, depending on the identified species (only for *Enterobacteriaceae*), performing a Multilocus Sequence Typing (MLST) classification, as well as a plasmid and virulence factor recognition. BaseSpace is a website developed by Illumina where registered users can easily store, analyze, and share genetic data (https://basespace.illumina.com). One of the benefits of BaseSpace is that users have access through their Illumina accounts, and the sequencing data from the sequencers, including iSeq100, are streamed in real-time over the Internet to BaseSpace at no additional cost. Once the sequencing data is in the BaseSpace Sequencing Hub, the user can access a limited set of free online BaseSpace apps for genomic and transcriptomic data analysis. However, most of the applications required for genomic raw data processing and bacteria species and ARG identification are not free. A yearly BaseSpace Sequence Hub Professional subscription costs $500 and includes 500 iCredits to go towards data storage or app usage. Additional credits can be purchased for ∼$1 each. The complete bioinformatic analysis using this platform costs about 7 iCredits or ∼ $7 per sample, of which 5 iCredits are for the Bacterial Analysis Pipeline alone. Additionally, BaseSpace failed to identify some of the ARGs found using other approaches, so it may not be the best option for diagnostic labs that require consistent, accurate results for all microorganisms.

### 2.4 M2gas a microbiome and physicocemical database

The *μ2gas* system was developed to support structured data storage and retrieval through a PostgreSQL database, a RESTful API, and a containerized deployment environment. The database schema was designed to manage interconnected datasets, with a central record table linking project metadata, plant specifications, chemical compositions, operational conditions, and output data. SQL scripts were used for schema initialization and data ingestion.

A Python-based API was implemented using Flask, enabling structured queries for data retrieval. The API architecture follows a modular design, separating request handling (rest_api.py), query execution (database_queries.py), and configuration management (config.py). The API exposes multiple endpoints for retrieving specific data subsets, supporting both raw database access and filtered queries.

To ensure portability and ease of deployment, the system was containerized using Docker. A set of Dockerfiles was developed for the database, API, and front-end, orchestrated via a docker-compose.yml configuration. The entire stack was deployed on an AWS EC2 instance, configured with Ubuntu Linux and supporting PostgreSQL, Flask, and NGINX for web service hosting.

**Table 1.**
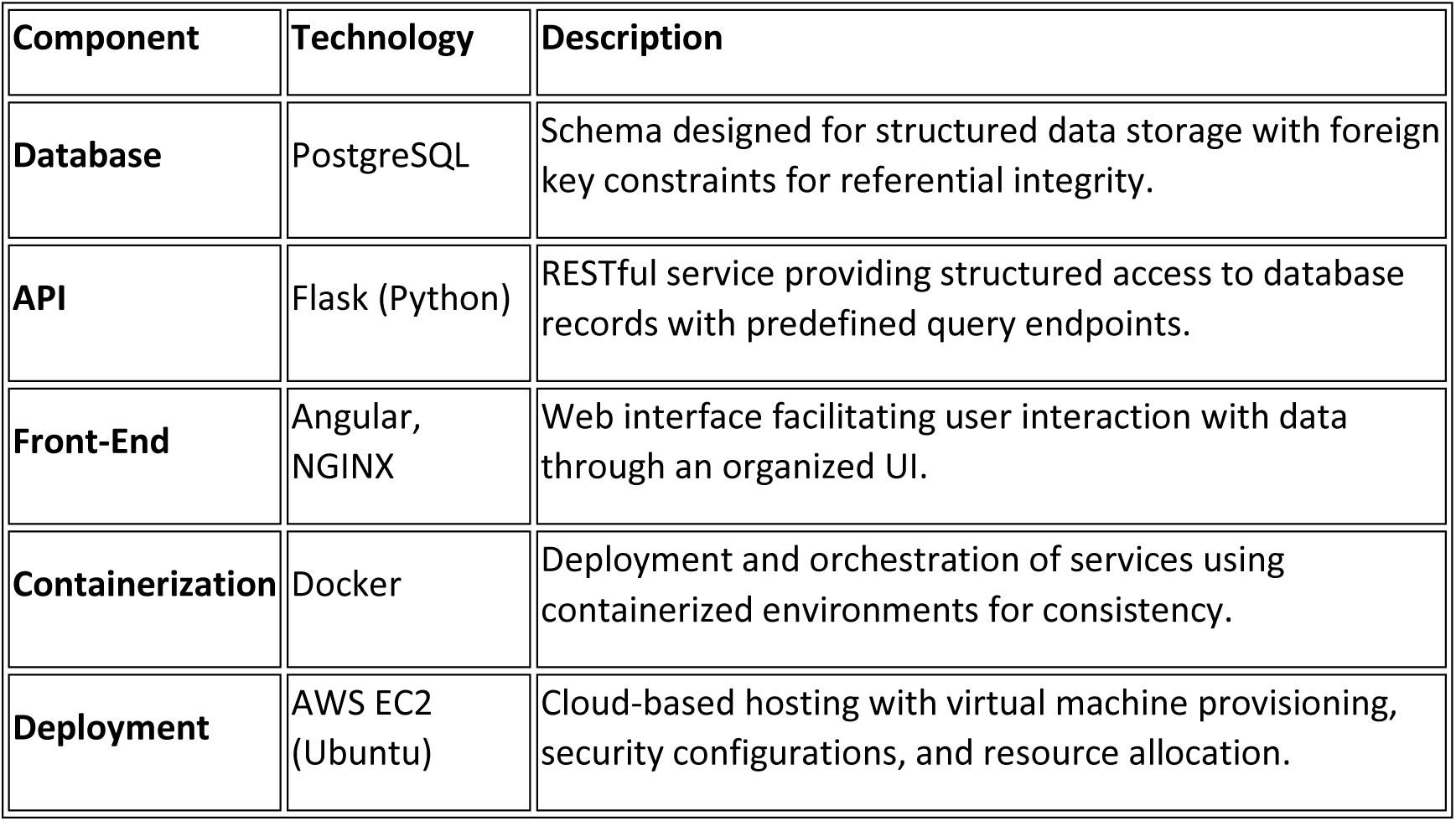
The following table summarizes the key components and their implementation details:

| Component | Technology | Description |
| --- | --- | --- |
| Database | PostgreSQL | Schema designed for structured data storage with foreign key constraints for referential integrity. |
| API | Flask (Python) | RESTful service providing structured access to database records with predefined query endpoints. |
| Front-End | Angular, NGINX | Web interface facilitating user interaction with data through an organized UI. |
| Containerization | Docker | Deployment and orchestration of services using containerized environments for consistency. |
| Deployment | AWS EC2 (Ubuntu) | Cloud-based hosting with virtual machine provisioning, security configurations, and resource allocation. |

## 3. Results

### 3.1 Microbial Community Dynamics and Reactor Stability

The microbial community dynamics and reactor stability were analyzed across different operational phases. The observed microbial shifts align with well-documented trends in anaerobic digestion (AD) systems, confirming the central role of core taxa in maintaining process stability and biogas production efficiency.

During steady-state operation, archaeal populations, particularly methanogenic genera such as Methanosarcina, maintained a stable presence (2–3%) (Fig. 1). However, during reactor destabilization, archaeal abundance dropped sharply (0.59% in July) (Fig. 2), correlating with the accumulation of volatile fatty acids (VFAs) and reduced methane yields. This pattern has been reported in similar studies, where methanogenic archaea were found to be highly sensitive to pH fluctuations and substrate overloads.

**Figure 1:**
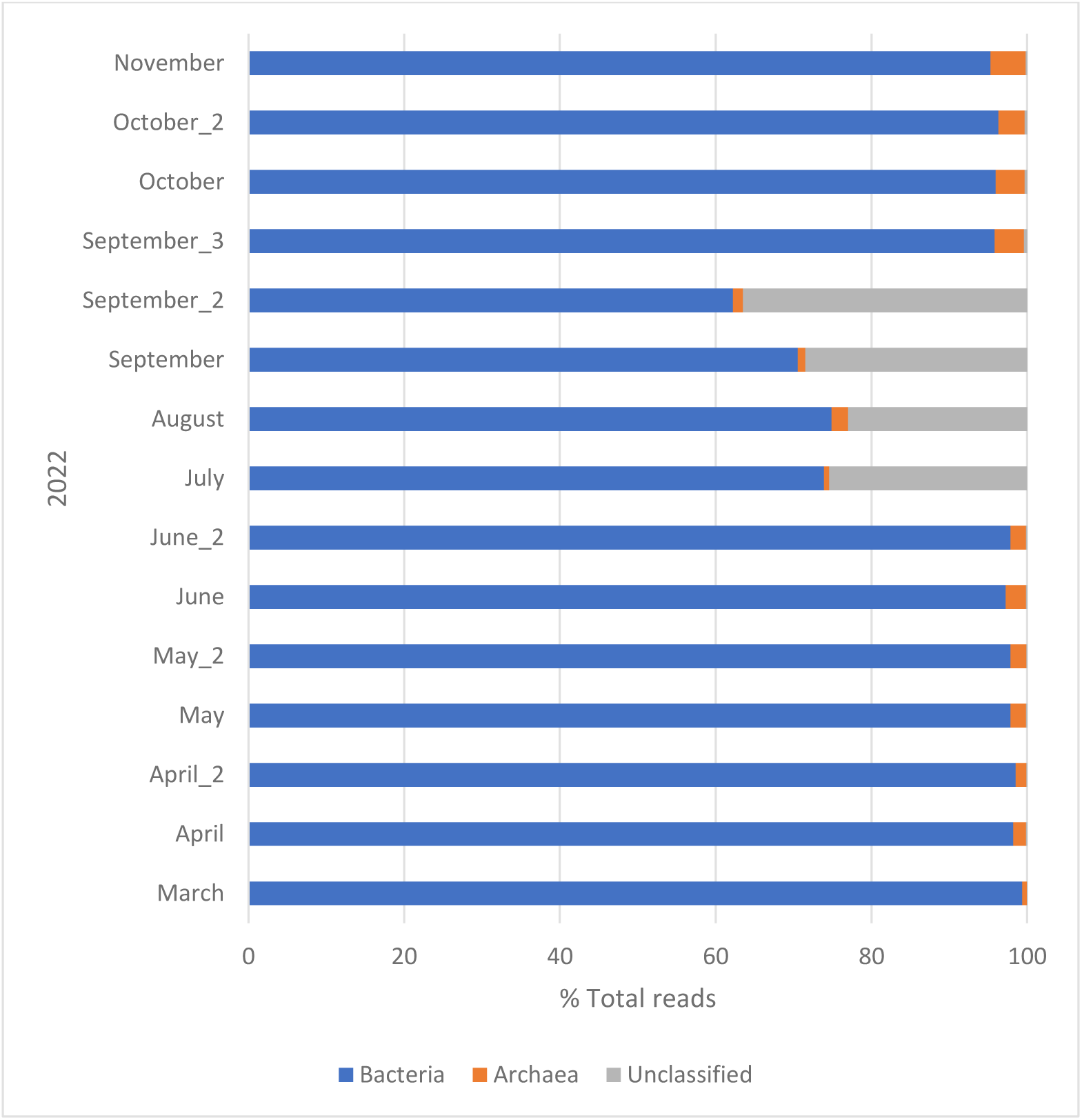
Microorganism identification rates at taxonomic kingdom level

**Figure 2:**
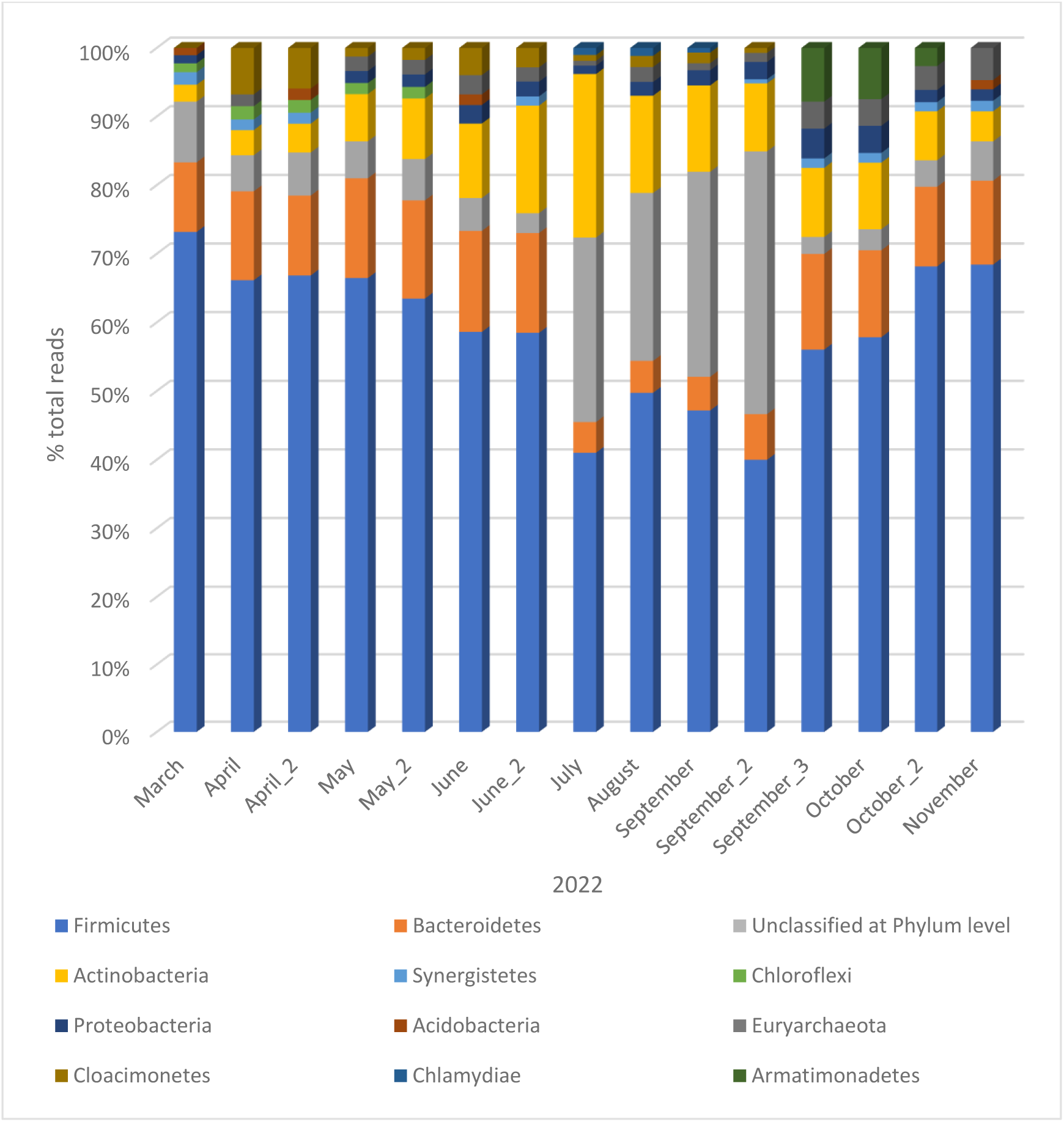
Microorganism identification rates at taxonomic phylum level

A significant decline in Firmicutes was observed during the destabilization phase, reaching its lowest levels in July (Fig. 3), only to recover gradually by November (Fig. 4). The decrease in Firmicutes, particularly Clostridia, aligns with previous findings that suggest their critical role in hydrolyzing complex organic matter into fermentable substrates for methanogens. The concomitant increase in Actinobacteria during reactor destabilization further supports the hypothesis that long hydraulic retention times and lower organic loads create favorable conditions for this phylum.

**Figure 3:**
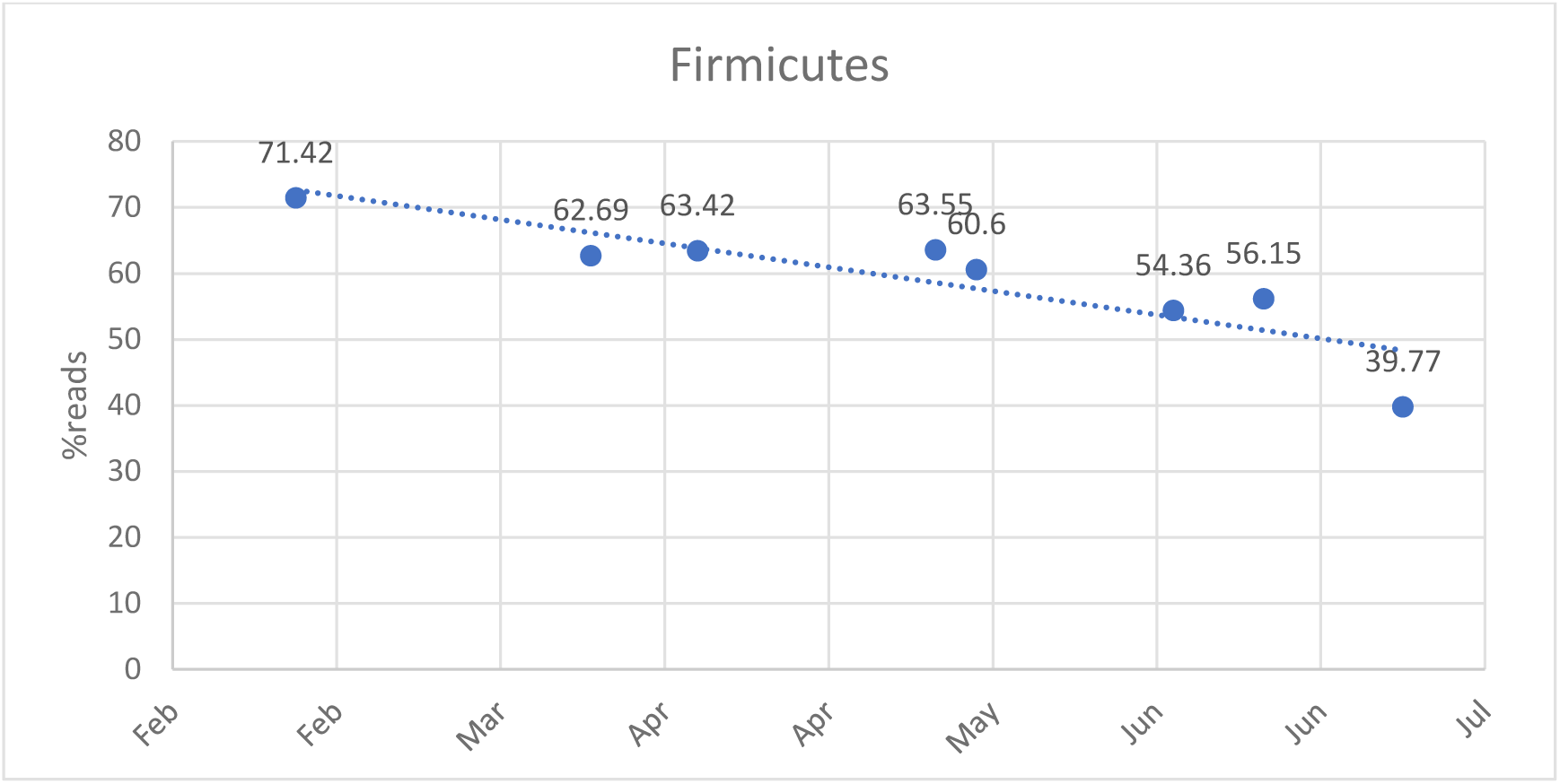
Gradual decrease of Firmicutes until July

**Figure 4:**
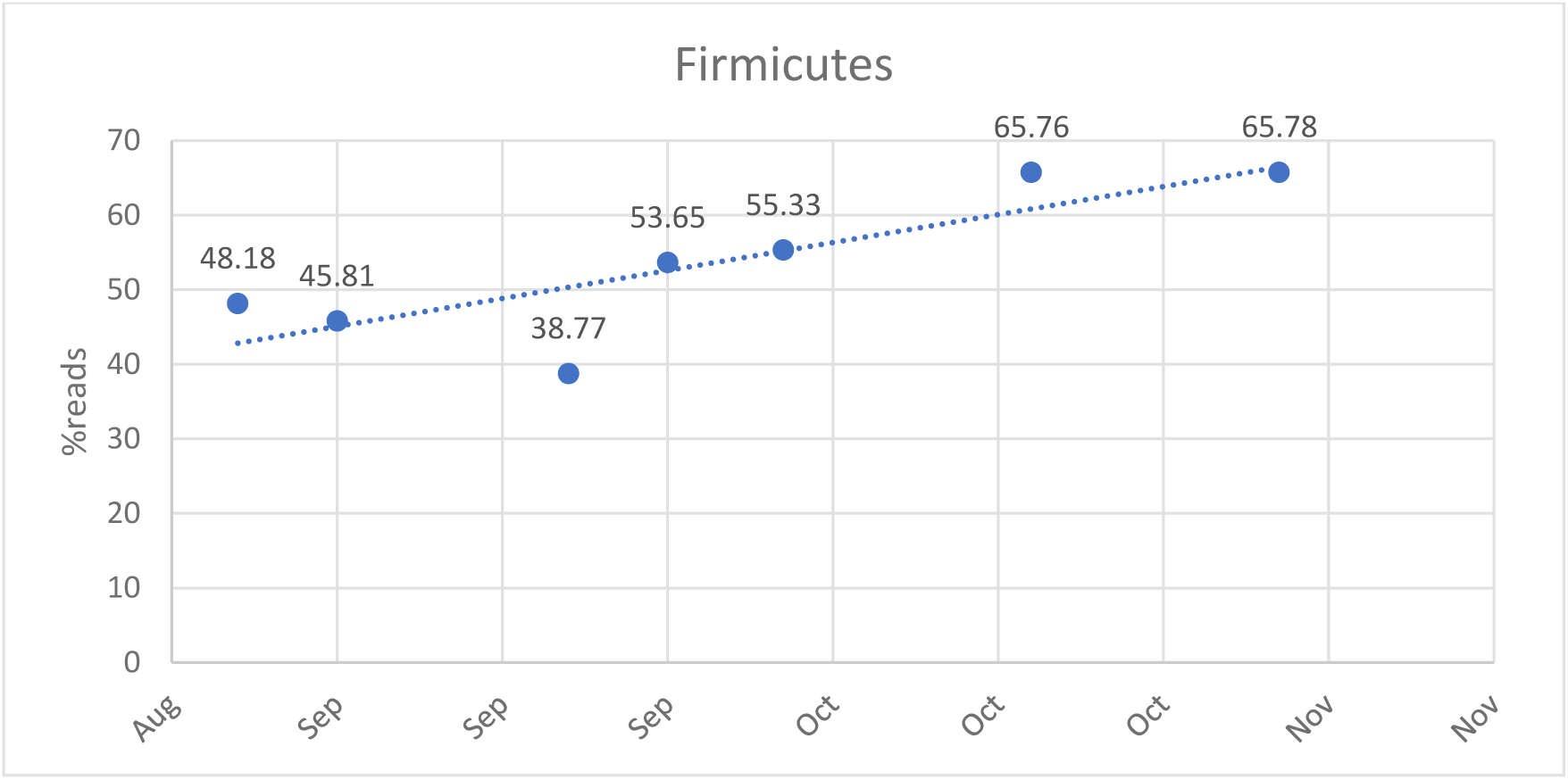
Gradual increase of Firmicutes until November

The sharp decline in Bacteroidetes (from 12% to 4.5%) (Fig. 5) suggests acidogenic inhibition, which aligns with the notion that Bacteroidetes thrive in conditions with low VFA accumulation and stable ammonia concentrations. The temporary increase in Armatimonadetes (7.5% in September) suggests its possible introduction via external feedstock inputs, further supporting reports that this genus is not a core community member in AD systems.

**Figure 5:**
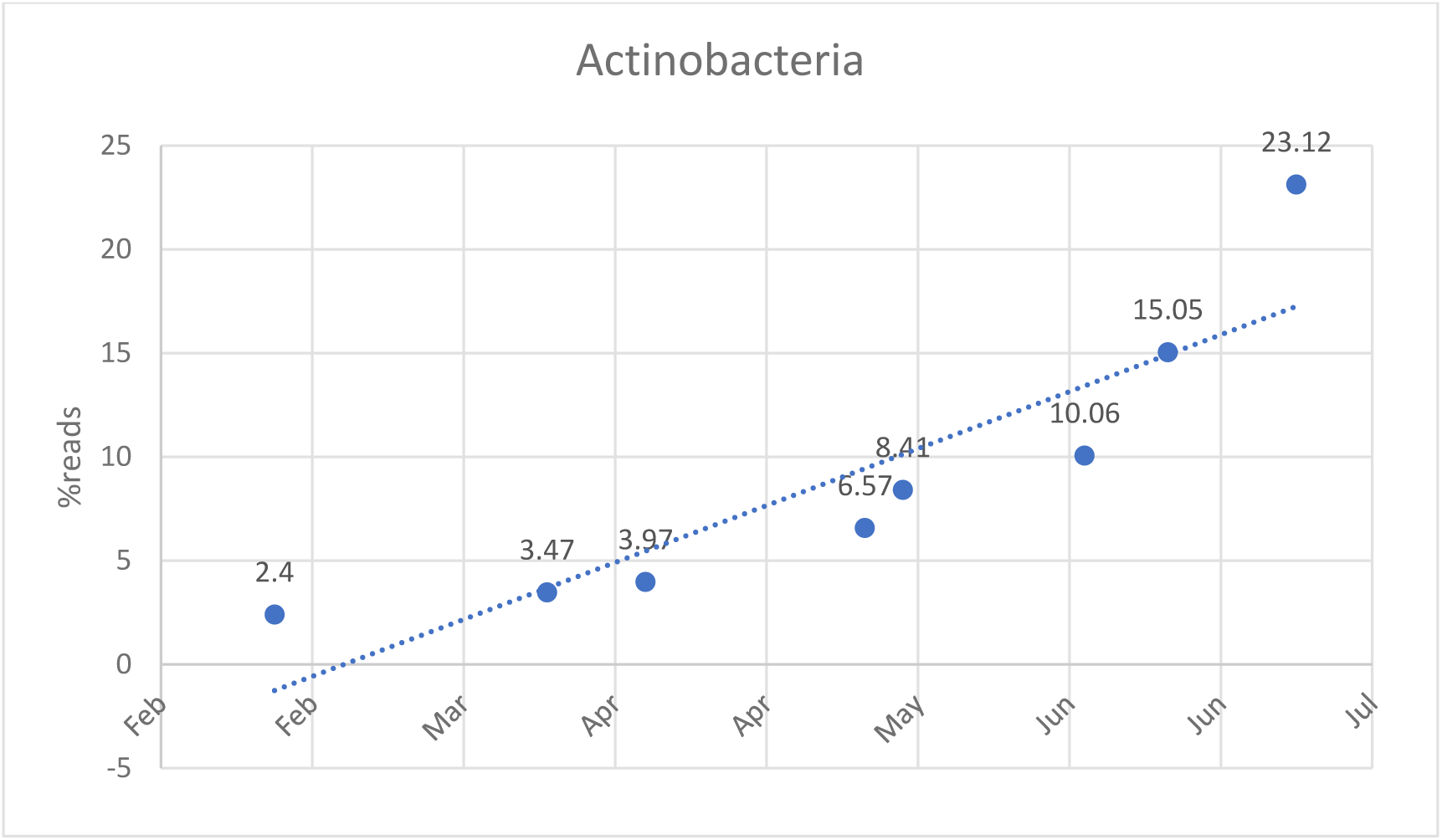
Exponential growth of the phylum Actinobacteria by July

### 3.2 Physicochemical Variations During Reactor Monitoring

The analysis of pH and temperature variations (Fig. S1) throughout the monitoring period demonstrated fluctuations that corresponded to key operational phases of the AD reactor. A decline in pH was observed during system destabilization, while temperature variations remained within the expected range, indicating that microbial imbalances rather than thermal stress were the primary contributors to reactor failure. The concentration of VFAs over time (Fig. S2) revealed an accumulation of these metabolites during the failure phase. This build-up coincided with a reduction in methanogenic activity, leading to an imbalance in microbial community structure. During reactor restoration, VFA concentrations gradually declined, aligning with the recovery of methanogenic archaea and improved process stability.

### 3.3 Feedstock Composition and Microbial Response

The monitoring of feedstock compositions (Fig. S3) highlighted key variations in input materials that influenced microbial dynamics. Seasonal shifts in feedstock sources led to compositional changes, which were reflected in microbial community adaptations. The introduction of certain organic substrates during restoration appeared to facilitate the recovery of key microbial taxa responsible for efficient anaerobic digestion.

The elemental analysis of feedstocks (Table S1) showed fluctuations in carbon, hydrogen, nitrogen, and oxygen content over time. These variations corresponded to shifts in microbial communities and reactor performance. Notably, periods with increased nitrogen content aligned with episodes of microbial stress, suggesting a potential inhibitory effect on certain microbial populations.

The detailed physicochemical characterization of different feedstocks (Table S2) demonstrated significant variations in moisture, ash content, volatile solids (VS), fat, protein, and theoretical gas yield. High moisture content in substrates such as liquid cow manure and whey correlated with lower VS percentages, which may have contributed to reduced biogas yields. Conversely, substrates with higher VS, such as solid chicken manure and glycerine, demonstrated greater theoretical gas yield potential, emphasizing the importance of feedstock selection in maintaining reactor efficiency.

### 3.4 Methane Production and Biogas Yield

Theoretical gas yield data (Table S2) provided insights into methane production efficiency across different feedstock types. The highest methane percentages were associated with glycerine and soap, indicating their potential as high-energy substrates for AD. In contrast, lower methane yields were observed for whey and silage, likely due to their lower organic content and higher moisture levels.

### 3.5 Application of the μ2Gas Database for Correlation Analysis

The observed correlations between microbial shifts, physicochemical variations, and reactor stability were made possible through the application of the μ2Gas database. This custom-built microbiome-oriented database facilitated the integration of sequencing data, operational parameters, and physicochemical conditions, enabling structured storage, retrieval, and correlation analysis. By leveraging its predictive analytics capabilities, the database allowed for early identification of destabilization events, aiding in targeted interventions to restore reactor function. The ability to track microbial community adaptations in response to operational changes highlights the value of μ2Gas in optimizing AD reactor performance and ensuring process resilience.

### 3.6 Implications for Reactor Optimization

The supplementary data emphasize the need for careful monitoring and optimization of feedstock composition, as variations in elemental and physicochemical properties significantly influence microbial community structure and overall reactor performance. The integration of microbiome-oriented data mining with physicochemical analysis provides a comprehensive approach to predicting and mitigating reactor instabilities. This study underscores the importance of microbiome-oriented monitoring for optimizing AD reactor stability. The observed microbial shifts highlight the necessity of real-time tracking tools to preemptively address destabilization events. Incorporating predictive bioinformatics models based on microbial abundance trends could enable earlier intervention and improved reactor resilience. Future research should focus on integrating machine learning techniques with microbiome monitoring to refine predictive models for reactor performance. Additionally, further metagenomic studies could provide functional insights into the metabolic pathways active during different operational states, enabling targeted strategies to enhance AD efficiency.Microbiome monitoring

A sharp decrease in the percentages was observed in July 2022. The percentages restored at the end of September 2022 where an increase of the percentage of the kingdom archaea was observed. The restoration of the rate of kingdom archaea appeared to be happening from August. In general, 2-3% of archaea microbial communities are found in anaerobic digestion reactors (REFERENCE). In the reactor under investigation, rates of 3.78% were detected at the end of September, 3.5% in October and 4.5% in November. Low rates were detected in March (0.59%) and July (0.68%).

A gradual decrease of Firmicutes was observed until July, but it was restored, also gradually, until November (Figures 3,4). The phylum Firmicutes mainly includes species that degrade cellulose (REFERENCE). Conversely, the phylum Actinobacteria showed a gradual increase until July and a decrease during the recovery period (Figures 5,6). Actinobacteriota species are favored at long hydraulic residence times and low organic loading of the reactor (REFERENCE).

**Figure 6:**
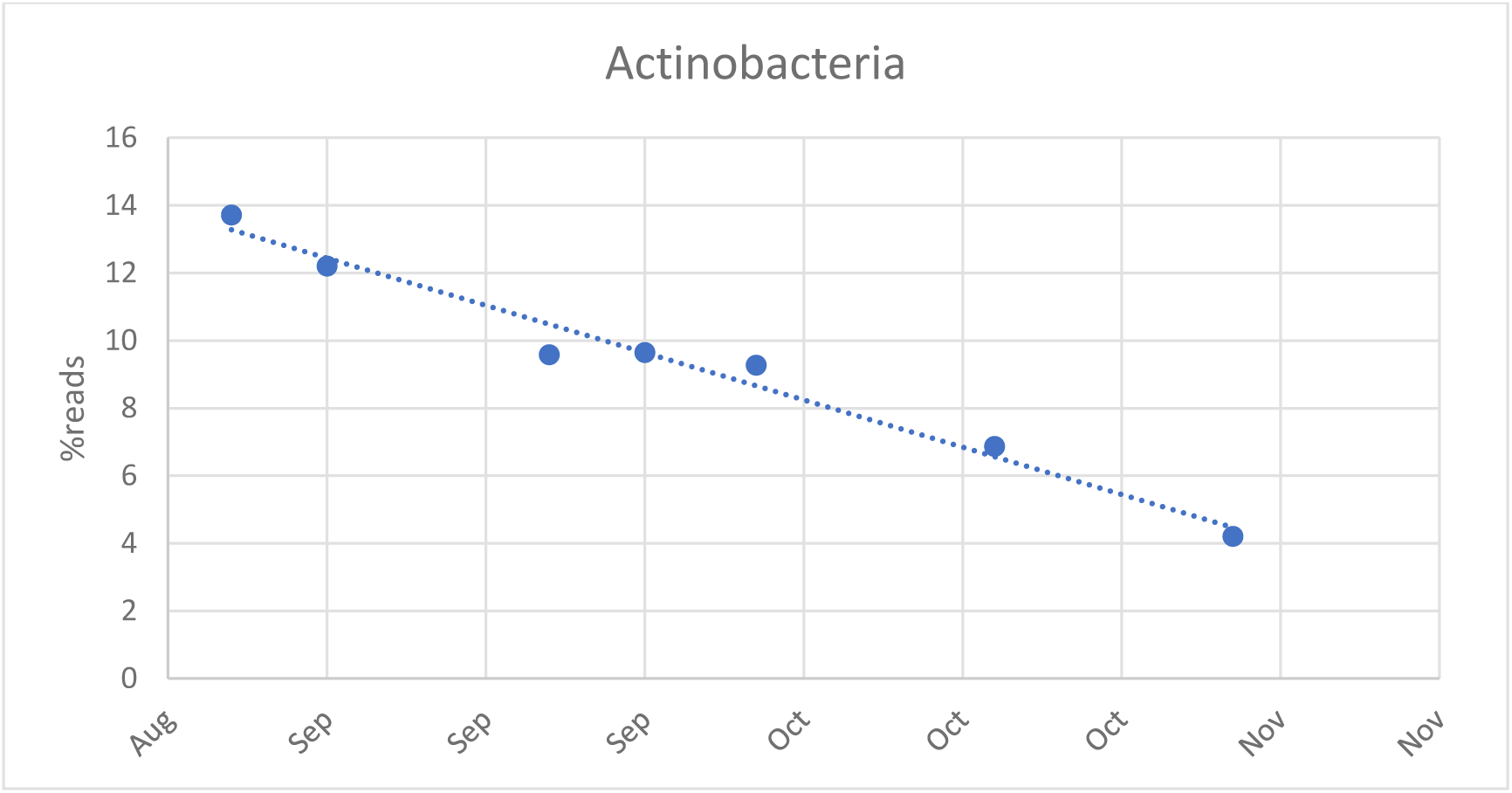
Linear decline of the phylum Actinobacteria until November

The percentage of Bacteroidetes seems to fluctuate around 12% under normal operating conditions, however it dropped to 4.5% in July and recovered at the end of September. This decrease suggests that acidogenesis also decreased during that period. Bacteroidetes have been reported to predominate in mesophilic reactors with low concentrations of volatile fatty acids (VFA), salinity and ammonia (REFERENCE).

It appears that 1.5% of the Synergistetes species is necessary for the proper operation of the reactor as this species was not found in the months of July until the end of September when it returned to 1.5%. Finally, the genus Armatimonadetes was found in high percentages at the end of September (7.5%) and October (7.15%). The genus Armatimonadetes is not a core genus of the reactor, so it is likely that it was introduced with new feed during the attempt to restore the unit.

The classes representing Firmicutes, Bacteroidetes and Actinobacteria following the above-mentioned changes are Clostridia, Bacteroidia and Actinobacteria, respectively. The class Clostridia, which includes species that break down carbohydrates and lipids, is reduced, as well as the class Bacteroidia, which has been associated with the breakdown of proteins and carbohydrates (REFERENCES).

The increase of Actinobacteria has been associated, recently, with the degradation of complex organic matter e.g. lignocellulosic substrates REFERENCE. Their high abundance during the destabilization period of the process is presumably related to an increase in the feed rates of the reactor in silage and chicken manure REFERENCE.

The phylum Bacilli, which has been associated with protein degradation, was not significantly altered. The relatively low percentages of the class Erysipelotrichia until the month of July (3.4%) suggest possible toxicity from ammonia accumulation (Figure 8). The class Negativicules appeared at 1.9-3.7% from late June to mid-September. There is no bibliographic data for the class Negativicules.

**Figure 7:**
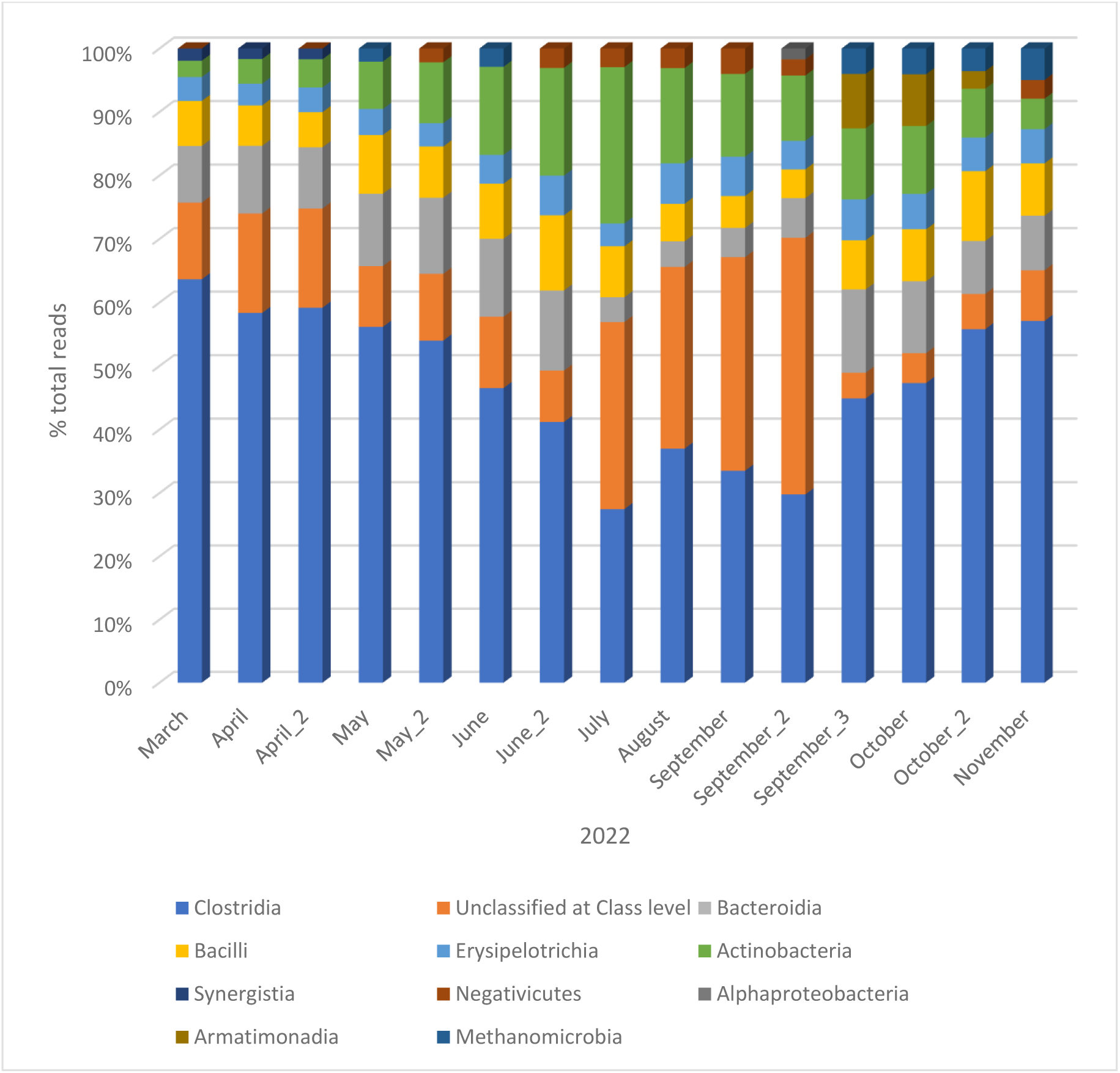
Microorganism identification rates at taxonomic class level

**Figure 8:**
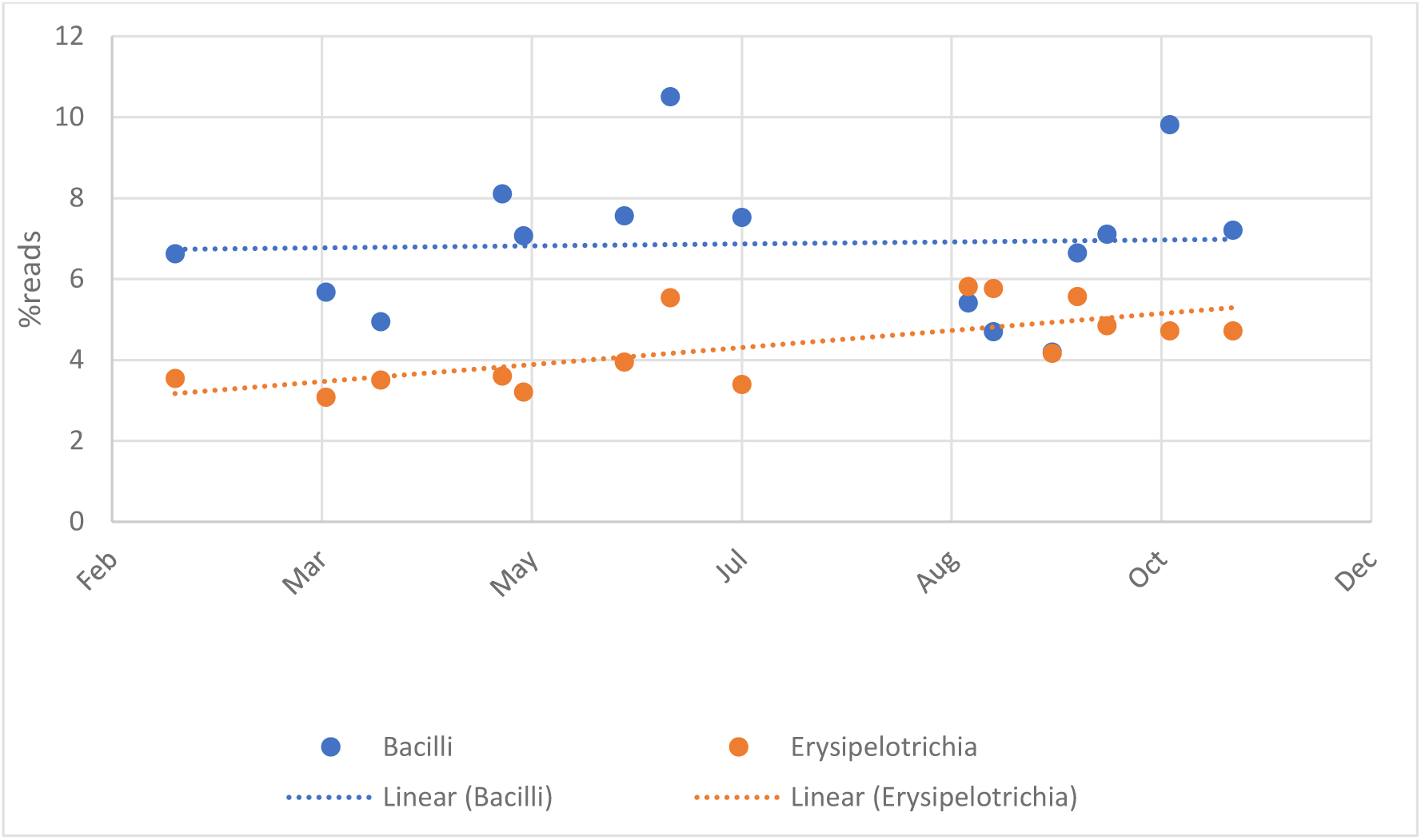
Change in abundance rates of the classes Bacilli and Erysipelotrichia during the monitoring interval of the unit.

Figure 10 shows the change in percentages of the four most abundant families over the reactor microbiome monitoring interval. It is observed that the families Peptostreptococcaceae and Erysipelotrichaceae were not affected by what caused the destabilization of the process but the families Ruminococcaceae and Clostridiales_Incertae_Sedis_XI appeared to decrease until July and increase thereafter. All four of these families play important roles in anaerobic digestion. The destabilization further reduced the abundance rates of the family Porphyromonadaceae, while, when the process was restored, increased rates of the family Lachnospiraceae (3.27-5.64%) were detected. These species have been associated with cellulose degradation to produce volatile oils acids (REFERENCE).

**Figure 10:**
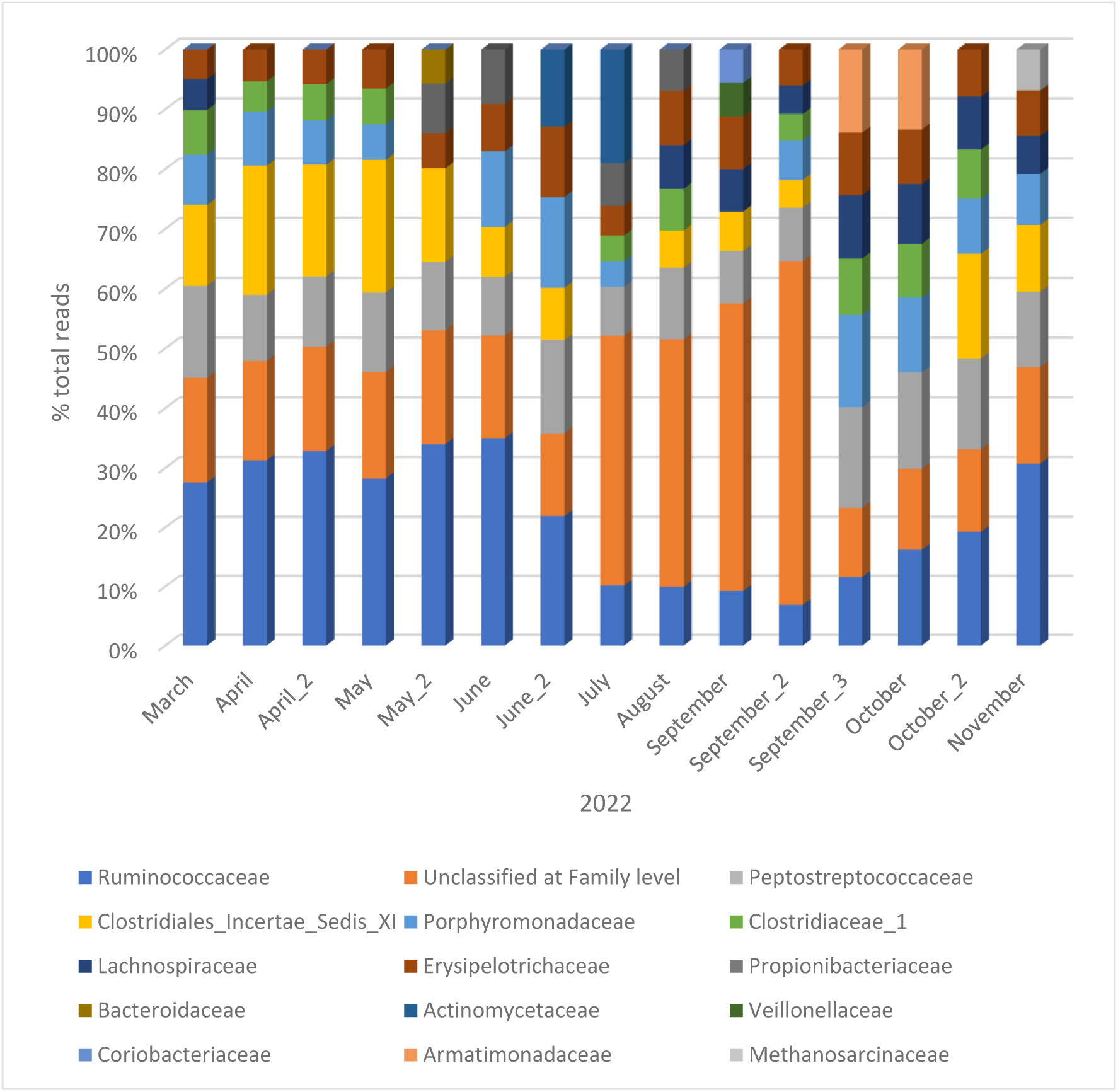
Microorganism identification rates at taxonomic family level

**Figure 11:**
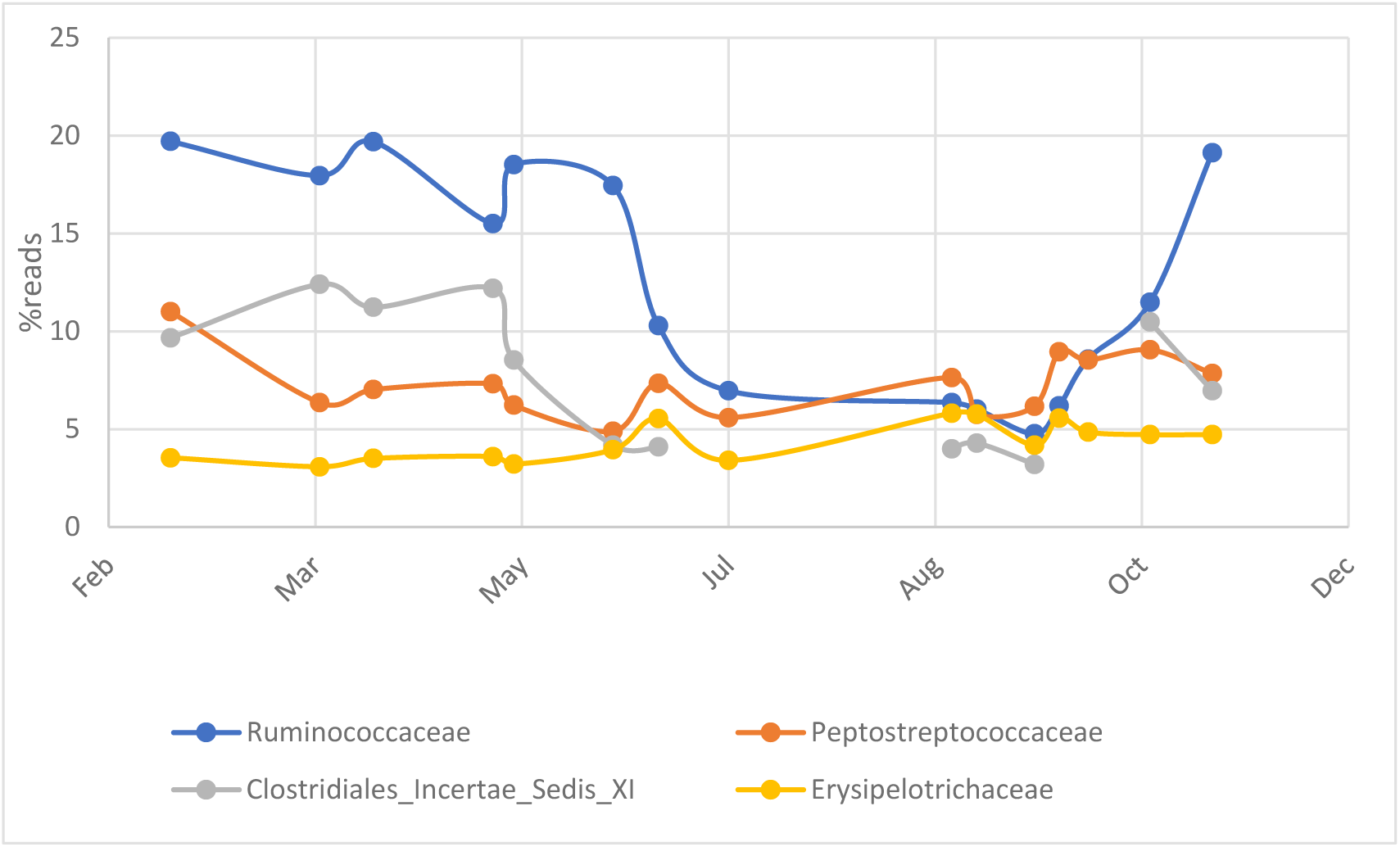
Variation of abundance rates of the four main families during the monitoring period of the unit.

Species of the family Ruminococcaceae are organotrophic bacteria, capable of degrading proteins, lipids and carbohydrates to produce volatile fatty acids (VFAs) and ammonia. They break down cellulose and produce hydrogen (H_2_) as one of the fermentation products. Some species produce propionate as a fermentation product (REFERENCE).

Species of the family Peptostreptococcaceae produce acid from glucose and other carbohydrates and are characterized by slow growth (REFERENCE). This slow growth favors their resistance over different organic loads and is legitimate for biogas production.

Finally, during the restoration phase of the unit, increased percentages of methanogenic archaea of the genus Methanosarcina were detected which are capable of acetotrophic and hydrogenotrophic methanogenesis depending on the balance of the chemical components of the reactor.

## 4. Discussion

The results of this study highlight the intricate relationships between microbial community composition and anaerobic digestion (AD) reactor performance across different operational phases. The observed microbial shifts align with well-documented trends in AD systems, confirming the central role of core taxa in maintaining process stability and biogas production efficiency. During steady-state operation, archaeal populations, particularly methanogenic genera such as Methanosarcina, maintained a stable presence (2–3%). This pattern is consistent with previous studies that emphasize the importance of a balanced archaea-to-bacteria ratio for efficient methanogenesis (Sundberg et al., 2013; Ziganshin et al., 2013; Maus et al., 2022). However, during reactor destabilization, archaea abundance dropped sharply to 0.59% in July, correlating with the accumulation of volatile fatty acids (VFAs) and reduced methane yields. This pattern is supported by prior research, which has reported that methanogenic archaea are highly sensitive to pH fluctuations and substrate overloads, leading to a decline in reactor performance (Chen et al., 2008; Schnürer & Jarvis, 2010; Wilkins et al., 2015).

A significant decline in Firmicutes was observed during the destabilization phase, reaching its lowest levels in July, only to recover gradually by November. Firmicutes, particularly the class Clostridia, play a crucial role in hydrolyzing complex organic matter into fermentable substrates for methanogens (Ahring et al., 2003; Callahan et al., 2016; Tian et al., 2020). Their decline suggests a loss of hydrolytic activity, which can lead to an accumulation of intermediate metabolites, further exacerbating reactor instability. Similar findings have been reported by Vanwonterghem et al. (2016), who demonstrated that shifts in Firmicutes abundance are key indicators of reactor health. Conversely, Actinobacteria exhibited an increase during the destabilization phase, likely due to prolonged hydraulic retention times and lower organic loading rates. This observation aligns with previous studies that have found Actinobacteria to be more prevalent under conditions of lower organic input and longer digestion times (Ma et al., 2021; Campanaro et al., 2020). The sharp decline in Bacteroidetes, dropping from 12% to 4.5% in July, indicates a disruption in acidogenic fermentation, which is essential for maintaining stable VFAs production (Wilkins et al., 2015). The temporary increase in Armatimonadetes (7.5% in September) suggests its possible introduction via external feedstock inputs, a hypothesis supported by studies indicating that Armatimonadetes are not core members of the AD microbiome but may proliferate under specific conditions (Ziganshin et al., 2013; Oyserman et al., 2021).

Comparative analyses with previous AD studies further support these findings. For instance, Sundberg et al. (2013) observed that archaea populations decline under increasing VFA concentrations, a trend that was replicated in this study. Similarly, Callahan et al. (2016) identified Firmicutes and Bacteroidetes as key phyla for reactor stability, with fluctuations in their abundance serving as early indicators of process disturbances. The shift from Firmicutes to Actinobacteria under stress mirrors findings by Vanwonterghem et al. (2016), emphasizing the adaptive potential of certain bacterial groups in response to changing organic loads. The presence of Synergistetes during stable operation but their absence during destabilization aligns with the findings of Ahring et al. (2003), who noted their role in syntrophic interactions with methanogens. The recovery of Lachnospiraceae and Ruminococcaceae post-destabilization supports previous reports on their function in hydrolytic and fermentative pathways crucial for biogas production (Jiang et al., 2021; Maus et al., 2022). These findings highlight the dynamic interactions between microbial taxa and operational conditions, reinforcing the necessity of continuous microbiome monitoring for optimal reactor performance.

All microbiome analyses were conducted using the custom-built microbiome-oriented database, μ2Gas, which integrated sequencing data, operational parameters, and physicochemical conditions. This system enabled efficient tracking of microbial shifts and facilitated comparative analysis with existing datasets. The integration of bioinformatics tools with microbiome monitoring represents a significant advancement in predictive analytics for AD optimization. Machine learning applications within the μ2Gas database successfully identified early warning indicators of failure, demonstrating strong potential for real-time process optimization and proactive intervention strategies. These findings underscore the importance of microbiome-oriented monitoring for optimizing AD reactor stability. The observed microbial shifts highlight the necessity of real-time tracking tools to preemptively address destabilization events. Incorporating predictive bioinformatics models based on microbial abundance trends could enable earlier intervention and improved reactor resilience. Future research should focus on integrating machine learning techniques with microbiome monitoring to refine predictive models for reactor performance. Additionally, further metagenomic studies could provide functional insights into the metabolic pathways active during different operational states, enabling targeted strategies to enhance AD efficiency.

### 4.1 Microbial Community Dynamics and Reactor Stability

During steady-state operation, archaeal populations, particularly methanogenic genera such as Methanosarcina, maintained a stable presence (2–3%)—a pattern consistent with previous studies indicating that a balanced archaea-to-bacteria ratio is crucial for efficient methanogenesis (Sundberg et al., 2013; Ziganshin et al., 2013; Maus et al., 2022). However, during reactor destabilization, archaea abundance dropped sharply (0.59% in July), correlating with the accumulation of volatile fatty acids (VFAs) and reduced methane yields. This pattern has been reported in similar studies, where methanogenic archaea were found to be highly sensitive to pH fluctuations and substrate overloads (Chen et al., 2008; Schnürer & Jarvis, 2010; Wilkins et al., 2015).

A significant decline in Firmicutes was observed during the destabilization phase, reaching its lowest levels in July, only to recover gradually by November. The decrease in Firmicutes, particularly Clostridia, aligns with previous findings that suggest their critical role in hydrolyzing complex organic matter into fermentable substrates for methanogens (Ahring et al., 2003; Callahan et al., 2016; Tian et al., 2020). The concomitant increase in Actinobacteria during reactor destabilization further supports the hypothesis that long hydraulic retention times and lower organic loads create favorable conditions for this phylum, as noted in prior research (Vanwonterghem et al., 2016; Yu et al., 2019).

The sharp decline in Bacteroidetes (from 12% to 4.5%) suggests acidogenic inhibition, which aligns with the notion that Bacteroidetes thrive in conditions with low VFA accumulation and stable ammonia concentrations (Ma et al., 2021; Campanaro et al., 2020). The temporary increase in Armatimonadetes (7.5% in September) suggests its possible introduction via external feedstock inputs, further supporting reports that this genus is not a core community member in AD systems (Ziganshin et al., 2013; Oyserman et al., 2021).

### 4.2 Comparative Analysis with Scientific Literature and Database-Driven Insights

The microbial shifts observed in this study are consistent with broader findings from anaerobic digestion research. Sundberg et al. (2013) identified similar trends in archaea fluctuations, emphasizing their vulnerability to operational disturbances. The decline of Methanosarcina during destabilization and its resurgence during recovery phases parallels findings by Ziganshin et al. (2013), reinforcing the notion that this genus exhibits resilience under fluctuating environmental conditions. Moreover, the observed fluctuations in Firmicutes and Bacteroidetes match findings by Callahan et al. (2016), who reported that these phyla are key indicators of reactor health and efficiency. The shift from Firmicutes to Actinobacteria under stress mirrors observations by Vanwonterghem et al. (2016), highlighting their adaptive potential in response to changing organic loads.

The presence of Synergistetes during stable operation but their absence during destabilization aligns with the findings of Ahring et al. (2003), who noted their role in syntrophic interactions with methanogens. The recovery of Lachnospiraceae and Ruminococcaceae post-destabilization supports previous reports on their function in hydrolytic and fermentative pathways crucial for biogas production (Jiang et al., 2021; Maus et al., 2022). All microbiome analyses were conducted using the custom-built microbiome-oriented database, which integrated sequencing data, operational parameters, and physicochemical conditions. The database allowed efficient tracking of microbial shifts and facilitated comparative analysis with existing datasets, confirming that microbial fluctuations correlated with reactor performance. This integration of microbiome data with real-time operational monitoring represents a significant step towards predictive analytics for anaerobic digestion optimization.

### 4.3 Implications for Reactor Optimization and Future Work

This study underscores the importance of microbiome-oriented monitoring for optimizing AD reactor stability. The observed microbial shifts highlight the necessity of real-time tracking tools to preemptively address destabilization events. Incorporating predictive bioinformatics models based on microbial abundance trends could enable earlier intervention and improved reactor resilience. Future research should focus on integrating machine learning techniques with microbiome monitoring to refine predictive models for reactor performance. Additionally, further metagenomic studies could provide functional insights into the metabolic pathways active during different operational states, enabling targeted strategies to enhance AD efficiency.

The findings underscore the intricate relationship between microbial community dynamics and reactor performance during different operational phases. The decline in Archaea and Firmicutes during destabilization aligns with the inhibition of methanogenesis and cellulose degradation, respectively. Conversely, the temporary increase in Actinobacteria and other less typical taxa, such as Armatimonadetes, suggests a shift toward stress-resilient microbial populations under adverse conditions. Restoration efforts, including feedstock adjustments and operational parameter optimization, successfully reinstated core microbial communities such as Archaea and Firmicutes, pivotal for reactor recovery. These observations are consistent with prior studies emphasizing the resilience and adaptability of microbial consortia in AD systems (Bolyen et al., 2019; Callahan et al., 2016). However, the transient dominance of non-core taxa highlights the need for further research into their roles and contributions during destabilization phases. Feedstock composition and seasonality emerged as critical factors influencing microbial dynamics, necessitating tailored strategies to mitigate variability’s impact on AD performance. The study also highlights the importance of integrating microbiome monitoring with physicochemical analysis to enhance predictive capabilities and reactor management. Future work should explore real-time monitoring and targeted interventions to maintain microbial stability and optimize biogas production.

## 5. Conclusion

This study underscores the intricate relationship between microbial community dynamics and anaerobic digestion (AD) reactor stability, highlighting the crucial role of microbiome-oriented monitoring in maintaining process efficiency. The observed microbial shifts, particularly the decline and subsequent recovery of methanogenic archaea, Firmicutes, and Bacteroidetes, emphasize the sensitivity of AD systems to environmental stressors and operational fluctuations. The integration of microbiome data with physicochemical analysis facilitated a comprehensive understanding of reactor performance, demonstrating the potential of data-driven approaches for early failure detection and predictive maintenance. The development and application of the **μ2Gas** database provided a structured framework for correlating microbial abundance patterns with reactor stability, reinforcing the utility of bioinformatics tools in optimizing biogas production. Machine learning models applied within this system successfully identified early warning indicators, paving the way for more proactive and adaptive reactor management strategies. These findings highlight the importance of leveraging microbiome data to improve AD resilience, enhance methane yields, and mitigate the impact of process disturbances. Future research should focus on real-time microbiome tracking, further refinement of predictive bioinformatics models, and the development of adaptive operational strategies to enhance reactor performance. Additionally, metagenomic analyses could provide deeper functional insights into microbial metabolic pathways, enabling targeted interventions to optimize AD efficiency. By integrating microbiome-oriented monitoring with machine learning-driven predictive analytics, this study contributes to advancing sustainable waste management and bioenergy production technologies.

## CRediT authorship contribution statement

**Manthos Panou:** Investigation, data curation, writing – original draft preparation, writing – review and editing. **Ioannis Kavakiotis**: Conceptualization, data curation, database creation.. **Vasiliki Tsioni:** Writing – review and editing, supervision, project administration. **Themistoklis Sfetsas:** Conceptualization, writing – review and editing, supervision, project administration, funding acquisition. All authors have read and agreed to the published version of the manuscript.

## Declaration of Competing Interest

The authors declare that there are no competing interests concerning the publication of this manuscript.

## Funding

This work was carried out as part of the project “μ2GAS” (Project code: KMP6-0143562) under the framework of the Action «Investment Plans of Innovation» of the Operational Program «Central Macedonia 2014-2020», that is co-funded by the European Regional Development Fund and Greece.

## Acknowledgements

The authors would like to acknowledge Ms. Afroditi Chioti for her work concerning the conceptualization and visualization of μ2Gas project.

## Notes

### Competing Interest Statement

The authors have declared no competing interest.

