## Supplementary Figures & Tables for "Microbiome-oriented data mining of operational monitoring of anaerobic digestion reactor during steady operation period, failure, and restoration"

Figure S1. **pH and Temperature Variations**

This figure presents the recorded variations in pH and temperature within the anaerobic digestion system over different time points.

|  | pH |
| --- | --- |
| 2022-02-07 | 7.9 |
| 2022-03-03 | 8 |
| 2022-03-11 | 7.9 |
| 2022-03-23 | 7.8 |
| 2022-03-31 | 8 |
| 2022-04-13 | 8.1 |
| 2022-04-15 | 8 |
| 2022-04-29 | 8 |
| 2022-05-03 | 8 |
| 2022-05-12 | 7.8 |
| 2022-05-17 | 7.9 |
| 2022-05-30 | 7.8 |
| 2022-06-10 | 8.1 |
| 2022-06-15 | 7.9 |
| 2022-06-20 | 8.2 |
| 2022-06-27 | 8.1 |
| 2022-07-08 | 7.9 |
| 2022-07-26 | 8 |
| 2022-08-17 | 8 |
| 2022-08-31 | 7.9 |
| 2022-09-06 | 8.2 |
| 2022-09-20 | 8.3 |
| 2022-09-26 | 8 |
| 2022-10-03 | 8.8 |
| 2022-10-18 | 8.1 |
| 2022-11-02 | 7.75 |
| 2022-11-11 | 7.8 |

FigureAC ratio

|  | FOS/TAC ratio |
| --- | --- |
| 2022-02-07 | 0.201 |
| 2022-03-03 | 0.211 |
| 2022-03-11 | 0.22 |
| 2022-03-23 | 0.173 |
| 2022-03-31 | 0.172 |
| 2022-04-13 | 0.163 |
| 2022-04-15 | 0.163 |
| 2022-04-29 | 0.15 |
| 2022-05-03 | 0.164 |
| 2022-05-12 | 0.166 |
| 2022-05-17 | 0.193 |
| 2022-05-30 | 0.16 |
| 2022-06-10 | 0.142 |
| 2022-06-15 | 0.154 |
| 2022-06-20 | 0.171 |
| 2022-06-27 | 0.181 |
| 2022-07-08 | 0.182 |
| 2022-07-26 | 0.186 |
| 2022-08-17 | 0.201 |
| 2022-08-31 | 0.256 |
| 2022-09-06 | 0.209 |
| 2022-09-20 | 0.191 |
| 2022-09-26 | 0.185 |
| 2022-10-03 | 0.126 |
| 2022-10-18 | 0.171 |
| 2022-11-02 | 0.225 |
| 2022-11-11 | 0.21 |

Figure S2. **Volatile Fatty Acids (VFA) Concentration Over Time**

This figure shows the variations in VFA concentrations, highlighting their influence on reactor stability and microbial performance.

VFA profile (ppm)

|  | Propionic acid | Isobutyric acid | Butyric acid | Isovaleric acid | Valeric acid | Isocaproic acid | Hexanoic acid | n-Heptanoic acid |
| --- | --- | --- | --- | --- | --- | --- | --- | --- |
| 2022-02-07 | 832 | 0 | 0 | 76 | 0 | 0 | 0 | 0 |
| 2022-03-03 | 453 | 32 | 0 | 28 | 0 | 0 | 0 | 0 |
| 2022-03-11 | 437 | 56 | 20 | 42 | 0 | 0 | 0 | 0 |
| 2022-03-23 | 0 | 0 | 0 | 0 | 0 | 0 | 0 | 0 |
| 2022-03-31 | 0 | 0 | 0 | 0 | 0 | 0 | 0 | 0 |
| 2022-04-13 | 0 | 0 | 0 | 289 | 0 | 0 | 0 | 0 |
| 2022-04-15 | 0 | 33 | 0 | 38 | 0 | 67 | 0 | 0 |
| 2022-04-29 | 32 | 0 | 0 | 56 | 0 | 0 | 0 | 0 |
| 2022-05-03 | 0 | 0 | 0 | 68 | 0 | 0 | 0 | 0 |
| 2022-05-12 | 42 | 0 | 0 | 0 | 0 | 0 | 0 | 0 |
| 2022-05-17 | 58 | 0 | 0 | 0 | 0 | 0 | 0 | 0 |
| 2022-05-30 | 58 | 0 | 0 | 0 | 21 | 0 | 0 | 0 |
| 2022-06-10 | 45 | 0 | 0 | 0 | 0 | 0 | 0 | 0 |
| 2022-06-15 | 59 | 0 | 0 | 0 | 0 | 0 | 0 | 0 |
| 2022-06-20 | 49 | 0 | 0 | 0 | 0 | 0 | 0 | 0 |
| 2022-06-27 | 179 | 0 | 0 | 0 | 0 | 0 | 0 | 0 |
| 2022-07-08 | 108 | 0 | 0 | 0 | 0 | 0 | 0 | 0 |
| 2022-07-26 | 105 | 0 | 0 | 0 | 0 | 0 | 0 | 0 |
| 2022-08-17 | 201 | 0 | 0 | 0 | 0 | 0 | 0 | 0 |
| 2022-08-31 | 293 | 17 | 39 | 0 | 44 | 0 | 0 | 46 |
| 2022-09-06 | 153 | 41 | 0 | 27 | 76 | 0 | 0 | 93 |
| 2022-09-20 | 242 | 0 | 26 | 0 | 0 | 0 | 0 | 0 |
| 2022-09-26 | 99 | 0 | 0 | 0 | 0 | 0 | 0 | 0 |
| 2022-10-03 | 0 | 0 | 0 | 0 | 0 | 0 | 0 | 0 |
| 2022-10-18 | 105 | 76 | 0 | 0 | 0 | 47 | 0 | 61 |
| 2022-11-02 | 153 | 0 | 0 | 0 | 0 | 0 | 0 | 0 |
| 2022-11-11 | 0 | 0 | 0 | 0 | 0 | 0 | 0 | 0 |

#### Figure S3. Feed Monitoring Trends

This figure presents the monitoring of feedstock compositions over time, highlighting key variations.

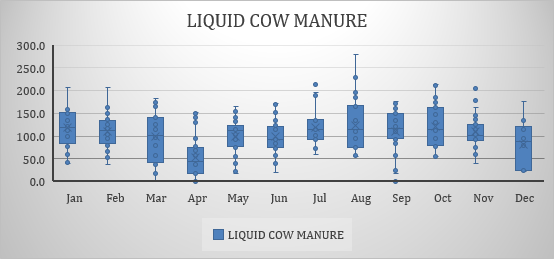

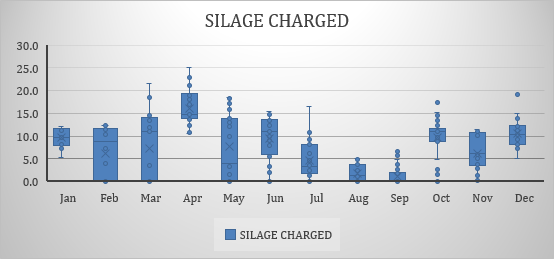

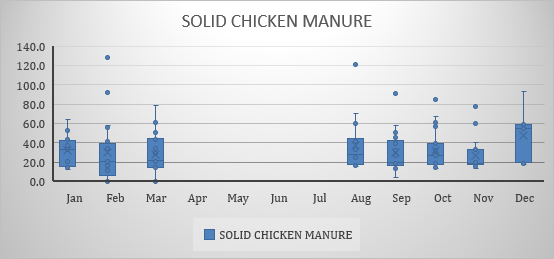

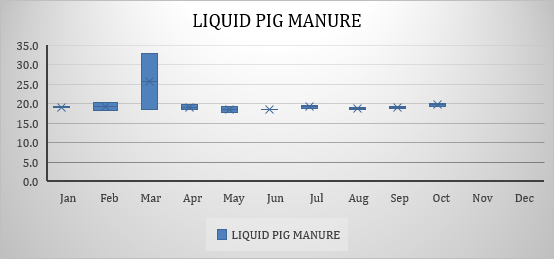

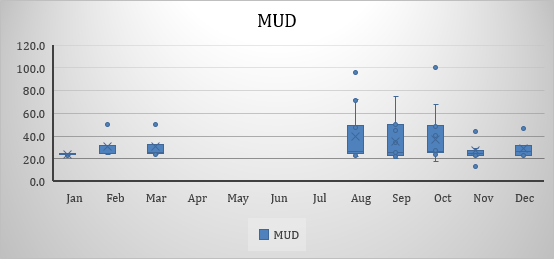

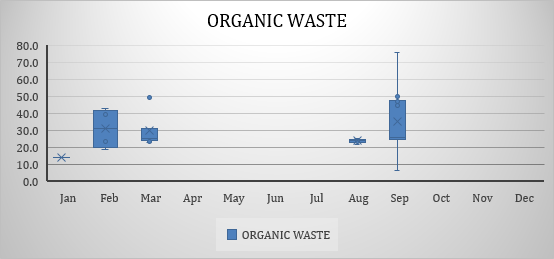

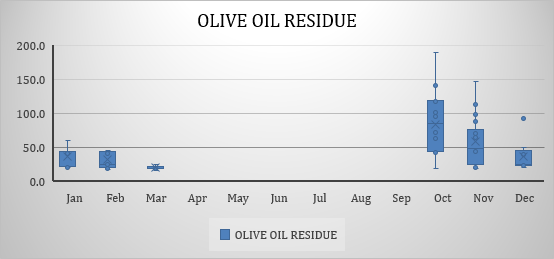

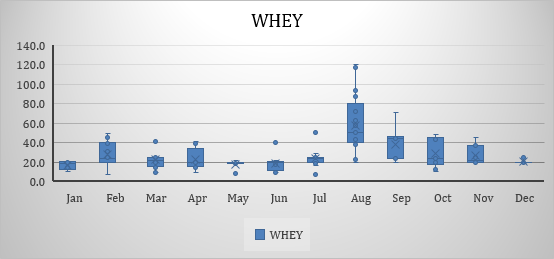

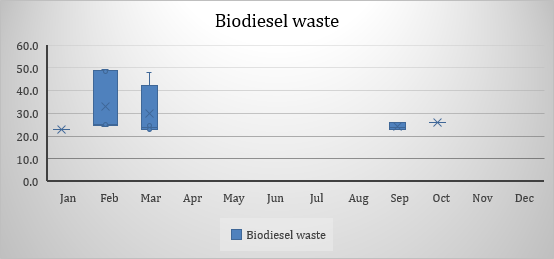

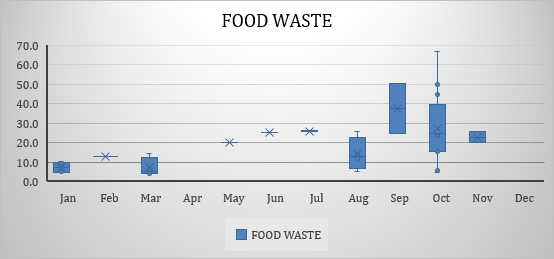

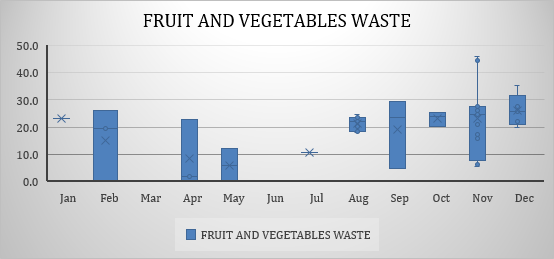

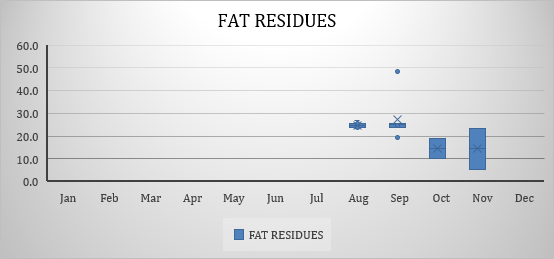

### Supplementary Tables

#### Table S1. Elemental Analysis (C, H, N, O₂ Composition Over Time)

Compositional data of feedstocks collected over multiple time points, indicating variations in elemental composition.

| Elemental analysis (%) | | |  |  |
| --- | --- | --- | --- | --- |
|  | C | H | N | O2 |
| 11/03/22 | 1.85 | 0.25 | 0.2 | 97.65 |
| 23/03/22 | 2.92 | 0.44 | 0.22 | 96.39 |
| 31/03/22 | 2.12 | 0.34 | 0.23 | 97.29 |
| 13/04/22 | 1.75 | 0.23 | 0.13 | 97.88 |
| 29/04/22 | 2.29 | 0.25 | 0.26 | 97.17 |
| 03/05/22 | 2.25 | 0.35 | 0.22 | 97.16 |
| 12/05/22 | 2.35 | 0.32 | 0.17 | 97.14 |
| 17/05/22 | 2.2 | 0.3 | 0.17 | 97.31 |
| 10/06/22 | 2.07 | 0.29 | 0.16 | 97.30 |
| 15/06/22 | 1.85 | 0.24 | 0.21 | 97.65 |
| 21/06/22 | 2.08 | 0.28 | 0.15 | 97.46 |
| 27/06/22 | 1.58 | 0.2 | 0.16 | 98.03 |
| 08/07/22 | 2.55 | 0.35 | 0.18 | 96.90 |
| 20/09/22 | 1.6 | 0.2 | 0.19 | 98.02 |
| 02/11/22 | 1.73 | 0.25 | 0.79 | 97.20 |

#### Table S2. Physicochemical Parameters of Different Feedstocks

Includes moisture, ash, volatile solids (VS), theoretical gas yield, and other relevant physicochemical properties.

| **Parameter** | Liquid cow manure | Solid chicken manure | Olive oil residue | Whey | Silage | Fruit & vegetable waste | Soap | Glycerine |
| --- | --- | --- | --- | --- | --- | --- | --- | --- |
| **Moisture (%)** | 93,16±2,8 (88,3-98) | 25±14,5 (13-52) | 91,2±7 (70,6-99,4) | 91,8±4,1 (83-97) | 69,6±9,4 (48,6-85) | 84,9±9,1 (64,3-99,5) | 88,2±7,2 (71,4-95,8) | 8,9±5,8 (1-22,5) |
| **Ash (%)** | 1,55±1,02 (0,57-3,4) | 15,8±4,14 (11,8-21) | 1±0,7  (0,1-2,5) | 1±0,9 (0,5-3,7) | 2,4±2,9 (0,8-13,5) | 0,7±1,2 (0,2-5,9) | 0,9±1,2  (0,34-5,2) | 3±1,9 (0,9-8,3) |
| **LOI (%)** | 98,16±1,02 (96,6-99,4) | 83,54±4,14 (78,9-88,2) | 98,8±0,7  (97,5-99,9) | 98,8±0,9 (96,3-99,5) | 96,8±2,9 (86,5-99,2) | 99±1,1 (94,2-99,8) | 98,7±1,2 (94,8-99,7) | 96,4±1,9 (91,8-99,1) |
| **VS (%)** | 5,81±2,96 (2-11,3) | 57,9±12,3 (38,3-68,7) | 4,9±6,6  (0,31-26,9) | 6,5±3,6 (3-15,2) | 25,5±8,8 (12,7-46,1) | 10,3±8,9 (0,52-33,9) | 8,7±6,2 (3,5-25,5) | 85,4±6,4 (71,1-98,1) |
| **Fat (%)** | 0,5±0,82 (0,18-2,46) | 2±0,9 (1,3-3,6) | 2,1±3,3  (0,2-14,1 | 0,32±1,16 (0,1-4,96) | 0,45±0,63 (0,02-2,72) | 0,57±1,63 (0,13-7,72) | 4,3±6,2 (3,5-24,6) | 1,4±30,5 (0,1-91,7) |
| **Proteins (%)** | 1,73±0,5 (0,18-2,5) | 18,1±5,4 (10,9-25,6) | 1±0,6  (0,4-2,9) | 0,9±0,9 (0,54-5,54) | 2,25±1,26 (0,6-6,2) | 1,64±1,6 (0,2-6,34) | -  (<0,5-1,71) | - |
| **Raw fiber (%)** | 1,35±0,37 (0,78-1,9) | 8,4±5,4 (3,9-17,4) | 0,77±2,85  (0,1-14) | - | 4,4±5,5 (0,82-19,7) | 2,8±3,8 (0,2-16,7) | -  (<0,3-0,4) | - |
| **Nitrogen free extracts (%DM**) | 11,4±14,2 (1,93-40,5) | 32,4±4,9 (24,8-38,2) | 15,9±18,8  (0,14-81,85) | 57,9±18,6 (9,4-87,5) | 50,4±21 (10,3-83,9) | 36,2±19,3 (2,2-84,4) | 16,5±27 (<0,5-84,4) | 51,9±32,8 (0,8-99) |
| **Theoretical gas yield (L/kg VS)** | 495±88 (382-629) | 451±87,4 (336-579) | 727±105  (474-935) | 639±54 (468-700) | 615±67,6 (446-685) | 582±51 (487-690) | 865±89 (634-988) | 834±122 (701-995) |
| **Methane (%)** | 58,8±5,7 (50,6-68,8) | 57,6±0,6 (56,7-58,4) | 58,5±5,9  (45,6-65,5) | 52,4±4,9 (42-65,9) | 51,3±5,9 (25,3-55) | 55±3,3 (49,5-65,7) | 56,7±5,7 (42,7-67,1) | 58,2±6,9 (50-66,9) |
| **pH** | 7±0,26 (6,6-7,3) | 8,1±0,34 (7,6-8,5) | 5,4±1,5  (4,2-12,5) | 4,3±0,8 (3,1-6,1) | 4,7±0,9 (3,6-6,6) | 4,3±1,2 (3,2-7,8) | 8,6±1,6 (5,8-11,9) | 4,4±3,5 (1,5-12,7) |
| **TS (%)** | 6,25±2,7 (2-11,7) | 70±14,5 (47,8-87) | 5,76±7 (0,61-29,4) | 7,2±4 (3-16,9) | 28±9,5 (15-51,4) | 10,6±9,3 (0,5-35,67) | 9,7±7,2 (4,2-28,6) | 88,8±5,8 (77,6-99) |

These supplementary materials provide additional data supporting the findings in the main text. They offer detailed insights into microbial community dynamics and reactor performance parameters.
